# CAR T cells targeting Nectin-4 safely overcome resistance to anti-Nectin-4 antibody-drug conjugate in solid tumors

**DOI:** 10.1101/2025.09.30.679440

**Authors:** Alexandrine Faivre, Léa Laban, Lorène Ferreira, Emmanuelle Josselin, Laurent Gorvel, Emilie Agavnian-Couquiaud, Anne Farina, Anne-Sophie Chrétien, Els Verhoeyen, Rémy Castellano, Jacques A. Nunès, Geoffrey Guittard, Marc Lopez, Daniel Olive

## Abstract

Chimeric antigen receptor T (CAR T) cells represent a promising therapeutic option for a variety of cancers, including solid tumors. Nectin-4 is a cell adhesion molecule expressed at different levels in many solid tumors, including breast and urothelial carcinoma (UC). Enfortumab vedotin (EV), an antibody-drug conjugate (ADC) against Nectin-4, has significantly improved survival in patients with metastatic UC. Skin adverse reactions are frequently observed due to Nectin-4 expression in the epidermis. Here, we developed second-generation CAR T cells against Nectin-4 (N4CART) with a scFv that does not recognize human Nectin-4 expressed in skin keratinocytes. To study the effects of N4CART cell therapy, we used preclinical models of cell-derived xenografts (CDX) and patient-derived xenograft (PDX) of breast cancer expressing moderate to high levels of Nectin-4. We showed a marked efficacy with induction of remissions. Interestingly, N4CART cells kill Nectin-4-positive breast, urothelial and colon tumor cells, which are resistant to EV. Thus, N4CART cells represent a valuable and safe therapy for the treatment of patients with Nectin-4 expressing tumors, including those that are resistant to EV. Finally, baboon-envelope pseudotyped lentiviral vectors (LV) outperformed VSVG-LVs for Nectin-4 CAR expressing in αβ T cells resulting in an efficient anti-tumoral response in PDX mice.

## Introduction

Chimeric antigen receptor (CAR) T cells against CD19 antigen (19CART cells) are able to provide very durable responses in patients with B-cell malignancies ^1-3^. However, CAR T cell efficacy against solid tumors remains elusive even though encouraging results have recently been reported ^4,5^. One of the difficulties is finding an ideal specific target antigen in solid tumors, which is of prime importance to improve and reach the efficacy observed in B-cell malignancies ^6^. Indeed, reduced antigen expression levels and low tumor specificity are criteria that can reduce the chance of successful CAR T therapy. There is a need to investigate and identify more accurate targets in solid tumors to optimize CAR T cell engineering.

PVRL4/Nectin-4, a type I transmembrane cell adhesion molecule, was previously described as a new marker, a tumor antigen and a target in breast and urothelial cancer ^7-9^. Nectin-4 is expressed both during fetal development, with expression declining in adult life, and as a tumor-associated antigen with pro-oncogenic properties in various carcinomas including breast cancer (BC) ^10-12 13,14^. Recently, enfortumab vedotin (EV), an antibody-drug conjugate (ADC) directed to Nectin-4, was FDA-approved in patients with locally advanced or metastatic urothelial cancer (la/mUC) ^15^. EV demonstrated an objective response rate of 41% in multiple pre-treated patients with la/mUC and 73% in association with pembrolizumab, and anti-PD1 immunotherapy ^16,17^. Skin toxicity is frequently observed due to Nectin-4 expression in keratinocytes, which limits the therapeutic index. These data validated Nectin-4 as a therapeutic target to treat la/mUC patients. However, resistance to EV has been described as also various alternatives to overcome this resistance ^18,19^.

An interesting alternative to the use of EV and ADC antibodies is the development of CAR-T cells expressing the accurate scFv. Previous study has described Nectin-4 CAR-T cells co-expressing IL-7 and CCL-19 that showed efficacy *in vitro* and in preclinical malignant cancer models such as lung, breast or bladder cancers. A phase I study NCT03932565, to examine the safety and feasibility of Nectin4-7.19 CAR-T cell infusion to patients, showed some major side effects such as haemorrhagic rash and rash desquamation ^20^.

Here, we developed Nectin-4-targeted chimeric antigen receptor-T (N4CART) cells with the aim of improving safety and anti-tumor efficacy *in vitro* and *in vivo* in various carcinomas. We also evaluated the efficacy of N4CART in EV-resistant models. Our results showed marked efficacy of N4CART cells in preclinical triple negative breast cancer (TNBC) PDX models and predicted minimal toxicity in clinical settings. This highlighted the N4CART cell potential to treat patients, even when refractory to treatment with anti-Nectin-4 ADC. Finally, we improved the transduction efficiency using baboon envelope pseudotyped lentiviral vectors to express Nectin-4 CAR in αβ T cells, which led to significant tumor regression in vivo in PDX mice.

## Results

### N4.78.6 mAb recognizes human and as well as mouse Nectin-4 antigen on tumor cells but does not bind keratinocytes

One of the important drawbacks of using either EV or already published CARN4 is the secondary effects triggered through the recognition of Nectin-4 notably in the skin. We thus decided to isolate a scFv that would ideally recognize Nectin-4 only on cancer cells and not on healthy tissues. Nectin-4 antibody selection was performed as previously described ^19^. The selected N4.78.6 antibody binds efficiently to Nectin-4 expressed on SUM190PT breast cancer cells and binds to mouse Nectin-4 (Fig1 A-B). IHC experiment on mouse skin tissue showed binding of N4.78.6 antibody to mouse skin (Fig 1C). We next compared by flow cytometry the binding of N4.78.6 to enfortumab vedotin (EV) on human keratinocyte (Fig 1D). Interestingly, both clones bind efficiently to Nectin-4 expressed on SUM190PT tumor cells, EV showing an increased affinity for Nectin-4 compared to N4.78.6 mAb (Fig 1D left). In contrast to EV-ADC, however, we could not detect the binding of N4.78.6 to human keratinocyte at any of the concentrations tested (Fig 1D right). This result was confirmed by immunohistochemistry (IHC) experiments to evaluate N4.78.6 binding on SUM190PT tumor tissue and on human and monkey skin tissues. Clearly, N4.78.6 binds Nectin-4 on SUM190PT tumor cells, but not on human and monkey skin. As control, we used the N4.72.1 that equally binds Nectin-4 expressed in tumor and normal skin tissues (Fig 1E). Together our data show that N4.78.6 efficiently recognizes Nectin-4 expressed on tumor cells but not on human and monkey skin.

**Fig. 1.**
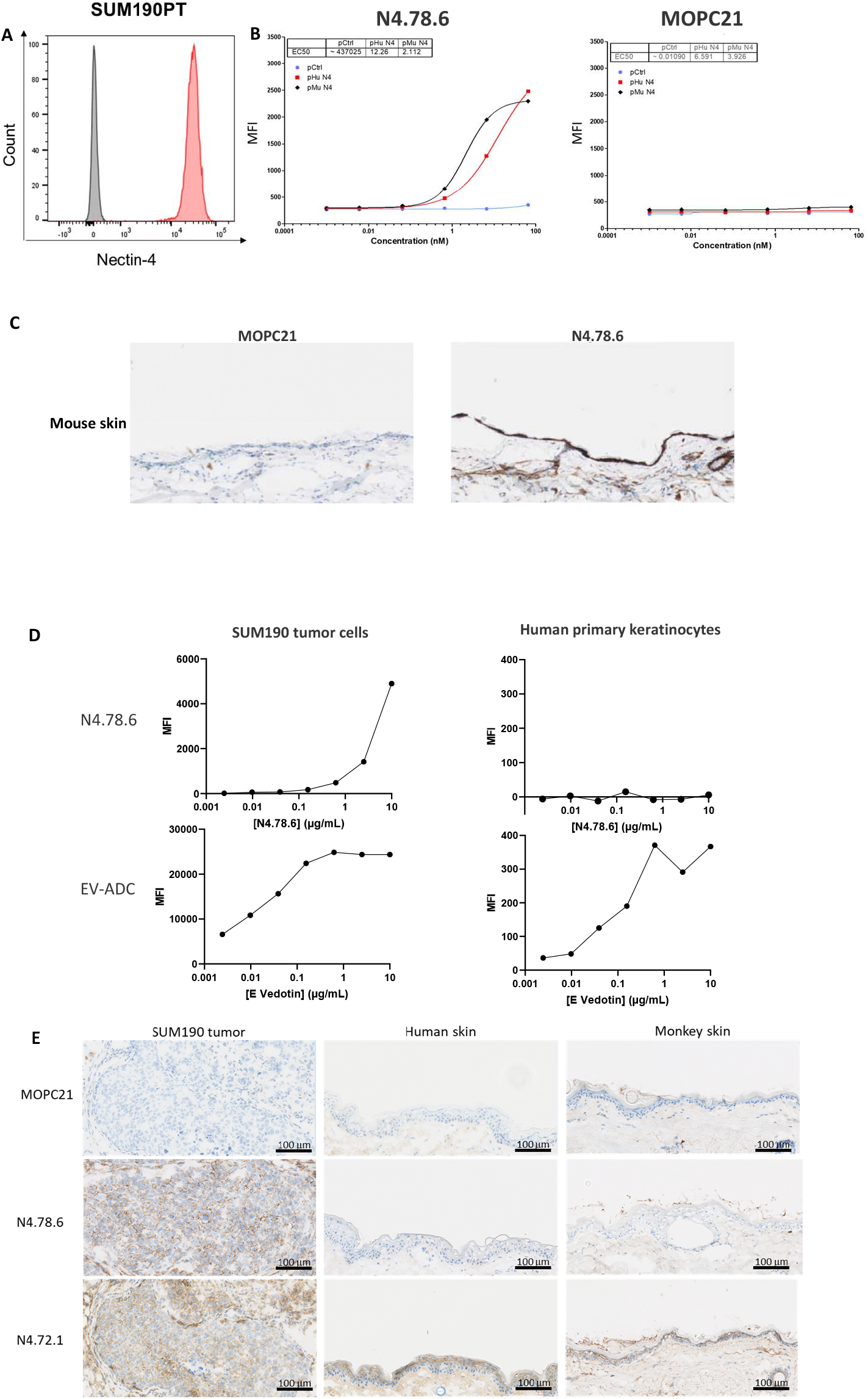
N4.78.6 mAb recognition properties. A. Nectin-4 expression on SUM190PT cell line with N4.78.6 antibody. B. (left) N4.78.6 binding on COS cells expressing human (red) or mouse (black) nectin-4. C. N4.78.6 staining of mouse skin. D. Comparison by FACS of N4.78.6 and EV-ADC binding on SUM190PT cells (left) and human primary keratinocytes (right). E. Comparison by IHC of N4.78.6 and N4.72.1 (skin positive control) staining of human and monkey skin. MOPC21: Mouse IgG1 isotype control. Data in panels A, B, C, E are representative of at least two experiments.

### A second-generation Nectin-4 CAR that targets efficiently Nectin-4 expressing tumor cells *in vitro*

To evaluate the efficacy of N4.78.6, we build two second-generation CARs with similar sequences except the scFv part either targeting Nectin-4 (N4CART) or CD19 (19CART). The variable region sequences of heavy (VH) and light chain (VL) of the anti-Nectin-4 N4.78.2 mAb were used to design the scFv connected with a G_4_S linker. We added to the construct the hinge and transmembrane part of CD8 and 4-1BB and CD3ζ as intracellular signaling domains. As showed previously N4.78.2 mAb was selected to bind human and murine Nectin-4 and to bind selectively to Nectin-4 expressed by human tumors and not by human skin keratinocytes (Fig1). To detect more easily N4CART cell expression using flow cytometry, we connected an eGFP sequence to the CAR constructs via a T2A ribosomal skipping sequence (Fig2.A). Prior to transduction, the peripheral blood mononuclear cells (PBMCs) were isolated from healthy donors and stimulated 48hrs with soluble anti-CD3, anti-CD28 and IL-2. Then, transduction with a VSV-G pseudotyped third generation lentivirus was performed at a MOI of 10 following a protocol established in the lab ^21^. Transduction reached around 40% of efficacy at day 3 post transduction as showed in cytometry by gating on cells double positive for GFP and the CAR, which was detected using a biotinylated N4 ^7^ (Fig-2B-C). The transduction did not change the CD4/CD8 ratio nor the T cell phenotype (suppl.Fig.1 A-B). Since Nectin-4 antigen is strongly expressed on SUM190PT TNBC cells, as shown by IHC experiments (Fig1E), we used this model in our subsequent *in vitro* experiments. 19CART cells or N4CART cells were co-cultured with SUM190PT at E:T ratio of 5:1 or with PMA and ionomycin (for maximal response) for 4h at 37°C. As expected, only CD4 and CD8 T cells transduced with Nectin-4-CAR expressed high levels of TNF-α and IFN-γ analyzed by FACS via intracellular staining, while 19CART cells did not. Both CAR T cells responded identically to PMA/ionomycin treatment (Fig2D). This was confirmed by detection of TNF-α and IFN-γ in the supernatants by ELISA (suppl.Fig.1 C). Similarly, N4CART cells showed an increased cytotoxicity compared to 19CART cells, as detected by caspase 3/7 labelling (Fig2E). Altogether our *in vitro* results showed functional expression of our CAR constructs and as expected an increased response of tumor-targeted N4CART cells compared to control 19CART cells.

**Fig. 2.**
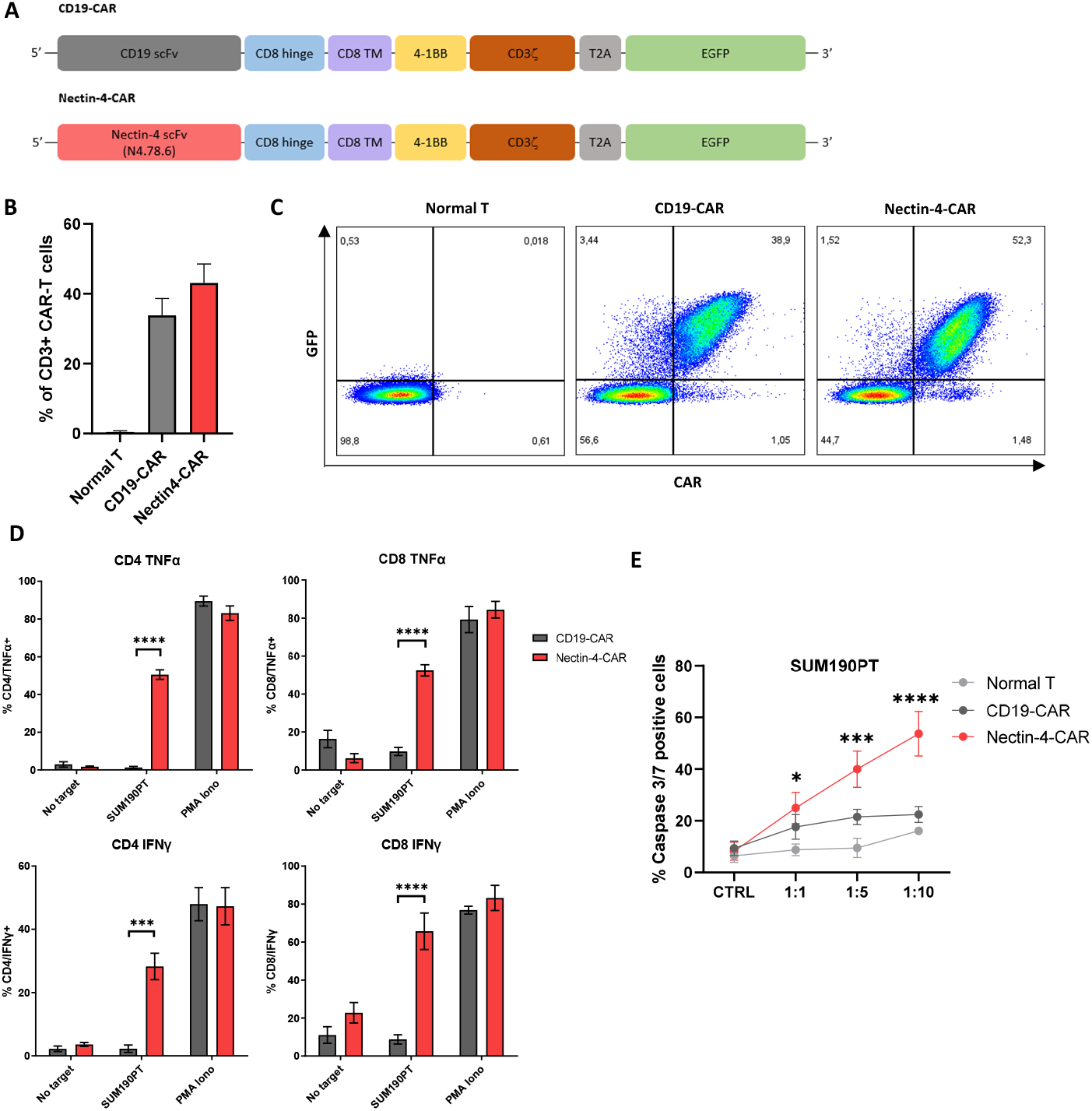
CARTN4 cells kill N4-positive tumor cell line and produce cytokines. A. Schematic diagrams showing the composition of the two CARs used in this study : CD19-CAR as control and Nectin-4-CAR. CARs was generated by fusing scFv to the costimulatory signaling domain of the 4-1BB and activating signaling domain of CD3z, a T2A ribosomal skipping sequence, and GFP was included for the detection of CAR-modified T cells. B. CD19 and Nectin-4 CAR expression reported by the GFP expression in activated human T cells upon VSV-G pseudotyped lentiviral transduction. n=4 healthy donors were examined in two independent experiments. C. Flow cytometry analysis of GFP and CAR expression reported by CD19 or Nectin-4 biotinylated recombinant protein. D. 4-hours TNFα and IFNγ production assay showing a significative production of cytokines by CARTN4 cells in contact with SUM190PT. CD4 and CD8 T-cells are permeabilized and stained with anti-TNFα and IFNγ antibodies. n=4 healthy donors were examined in two independent experiments. E. 4-hrs caspase 3/7 cytotoxicity assay indicating a higher tumor killing CARTN4 cells against target cells. n=4 healthy donors were examined in two independent experiments. Data in panels B, E, F are presented as Mean +/- SEM. P-values were calculated using a 2way ANOVA. Source data are provided in the Source Data file.

### N4CART cells efficiently inhibit tumor progression *in vivo*

We next evaluated the efficiency of N4CART cells *in vivo*. NSG mice were inoculated with 5 × 10^5^ SUM190PT cells in mammary fat pads of both flanks. When tumors reached an average volume of 80-100 mm^3^, mice were randomized then received an adoptive transfer of 5 × 10^6^ transduced CAR-T cells (Fig2A) that have been *ex vivo* amplified for 7 days (Fig 3A). Tumor growth was monitored every 3-4 days post-injection and the tumor volume was assessed (Fig 3B). As expected, a major decrease in tumor growth was detected in tumor-bearing mice injected with N4CART cells compared to 19CART cells. We collected and harvested tumors at day 10 post-injection to immunophenotype the lymphocytes. We observed a significant enrichment of CAR+ CD3+ cells in the tumors injected with N4CART cells compared to 19CART cells, predominantly due to the increase in CD4+ CAR+ T cells (Fig 3C; suppl. Fig 2A). We then studied the phenotype of CAR infiltrated lymphocytes (CILs). Using CD45RA and CD27 labelling we showed an increase in the effector memory population (Tem; CD45RA-CD27-) and a decrease in terminally differentiated effector memory cells re-expressing CD45RA (Temra; CD45RA+ CD27-) in CILs from N4CART as compared to 19CART that is observed either for CD4+ or CD8+ expressing CAR-T cells (Fig3-D-E; suppl. Fig 2B). CD4+ and CD8+ infiltrated N4CART cells express a lower amount of PD-1 and Tim-3 inhibiting receptors at their surface when compared to infiltrated 19CART cells (Fig.3F-G; suppl.Fig 2C). Of note, we did not observe any weight loss of the different injected mice (suppl.Fig 2D). We also compared the phenotype status of tumor infiltrated CAR+ versus CAR-CD3+ T cells from the same tumor (suppl.Fig 3A-C). Consistent with the FACS data, we observed an increased representation of either CD4+ or CD8+ CAR+ T cells compared to the CAR-T cells. Additionally, CAR+ T cells tended to be less EMRA and less exhausted compared to CAR-infiltrated T cells (suppl.Fig 3A-C). ELISA was performed on both tumor supernatants revealing an important production of TNF-α and IFN-γ when N4CART cells infiltrated the tumor compared to 19CART cells (Fig 3H). We finally performed an immunohistochemistry (IHC) labelling for GFP and CD3 T cells on the tumor sections (Fig 3I). Consistent with the cytometry results, we observed a marked intra-tumoral infiltration of GFP+ CD3+ CAR-T cells in the N4CART cell condition as compared to 19CART cells.

**Fig. 3.**
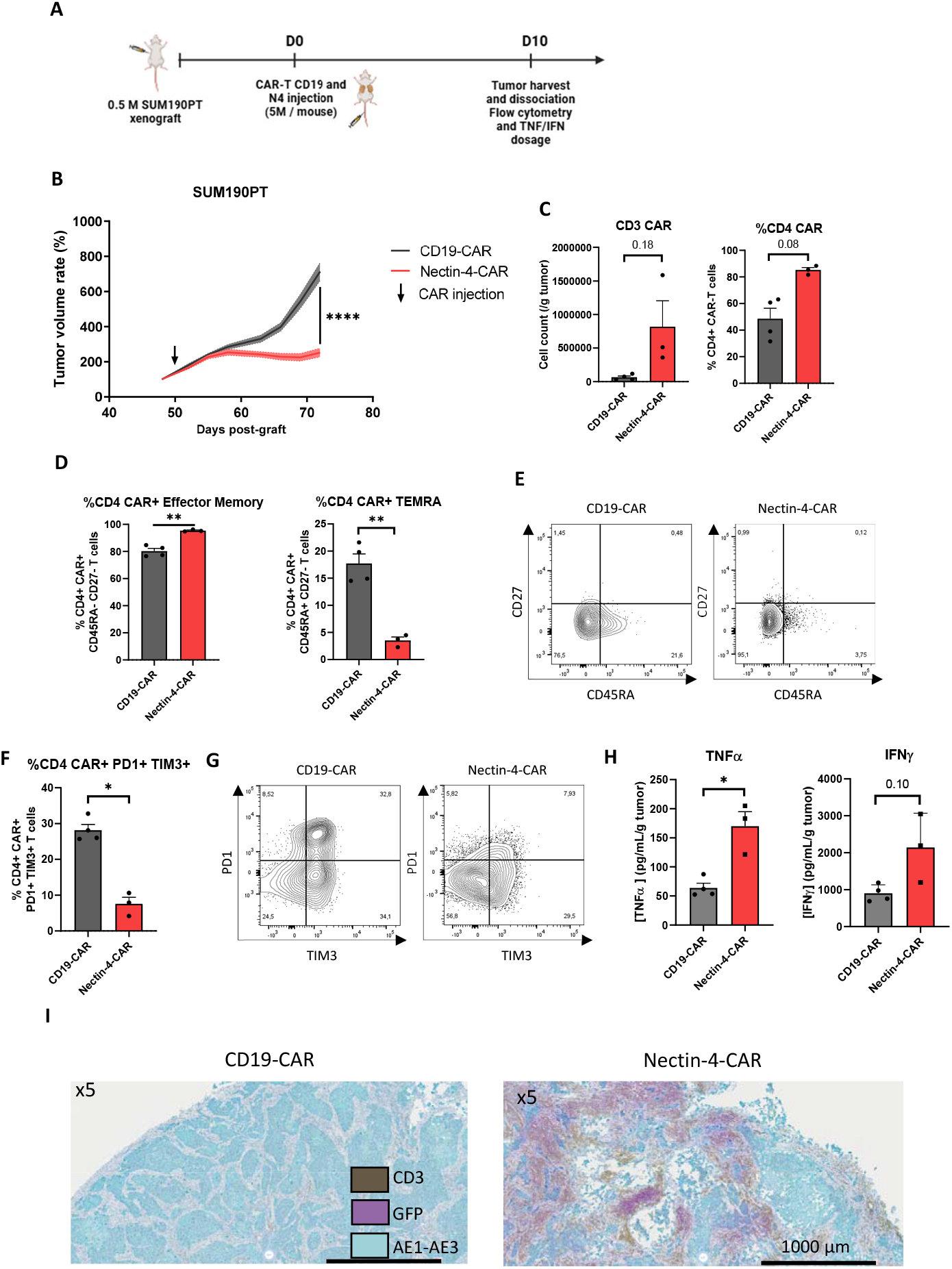
Mice “one-shot” treatment with CARTN4 cells leads to tumor growth control in human tumor xenograft models. A. Schematic illustration of the protocol. B. SUM190PT tumor growth control by a single injection of CARTN4 cells *in vivo*. SUM190PT cells were orthotopically injected into the mammary fat pad of female NSG mice. Mice received a single i.v. injection of CART19 as vehicle control or CARTN4 cells. n = 5 CD19-CAR and n = 5 Nectin-4-CAR animals with 2 tumors each were examined in one experiment. C. CD3+GFP+ absolute count per gram of tumor and CD4 rate are higher in Nectin-4 tumor. Tumors were harvested and dissociated 10 days after CAR-T cells treatment. D. Effector memory and EMRA T-cells rate in CD4 T-cells in tumor at day 10. E. Flow cytometry results illustrating the higher frequency of TEMRA in CD19-CAR-T cells and the higher frequency of effector memory T-cells in Nectin-4-CAR-T cells. F. PD1 and TIM3 double positive T-cells rate in CD4 T-cells in tumors at day 10. G. Flow cytometry results illustrating the higher frequency of PD1 and TIM3 double positive T-cells in CD19-CAR-T cells showing a more exhausted phenotype. H. ELISA assay dosing TNFα and IFNγ in tumor dissociation supernatants revealing higher concentrations of both cytokines in CARTN4 / tumors conditions. I. IHC on mouse SUM190PT tumor 10 days after CAR treatment showing a massive CARTN4 cell infiltration. Each point represents one tumor. 4 tumors were harvested from 2 mice per condition. Data in panels B, C, D, F and H are presented as Mean +/- SEM. P-values were calculated using a 2way ANOVA in panel B and using a t-test in panel C, D, F and H. Source data are provided in the Source Data file.

### Complete tumor regression is achieved upon CAR-T Double injection

Since we observed a marked yet not complete regression with a single injection of N4CARTcells, we added another injection 10 days after the initial one (Fig 4A). The second injection clearly abrogated tumor progression and we could observe 9/10 complete responses using this new approach (Fig 4B). We next studied T cell infiltrations at day 3, 10, 12 and 20 post-initial first injection. As expected, N4CART cells infiltrated more efficiently the tumor at any of these time-points compared to 19CART cells (Fig 4C). Interestingly, the second injection boosted even more CAR-T cell infiltration into the tumors, which peaked 2 days after injection to decrease again at day 20 correlating with remarkable tumor regression (Day 45 after the tumor cell engrafting). We also observed that the second injection allowed a change in the percentage balance of CD4 versus CD8+ CAR+ T cells at day 20 in infiltrated N4CART cells (Fig 4D; Suppl. 4A). As observed previously for the single injection (Fig 3), we detected an enrichment in CD4+ and CD8+ TEM and a decrease in TEMRA at day 10 and 12 of infiltrated N4CART cells compared to 19CART. A very similar T cell phenotype was detected at day 20 for the two different CAR T cells, possibly due to rapid tumor collapse leading to N4CAR T cell regression (Fig 4 E-F, suppl. Fig 4B). We observed the same for T cell exhaustion markers where N4CART cells showed less PD1+ TIM3+ expression at day 10 and 12 compared to 19CART cells, while both CAR T cell types reached a similar expression profile at day 20 (Fig 4G-H, suppl. Fig 4C). No obvious toxicity was detected since we did not observe any weight loss of the different injected mice, even upon two CAR T cell injections (suppl. Fig 4D). Similar to a single CAR T cell injection, an increased recruitment of either CD4+ or CD8+ CAR+ T cells compared to the CAR-T cells was observed and CAR+ T cells tended to be less EMRA and less exhausted compared to CAR-infiltrated T cells (suppl. Fig 5A-B). Cell-free supernatants were harvested and an increase in the expression of TNF-α and INF-γ by ELISA was observed, reaching a maximum at day 20 (Fig. 4I). This last observation confirmed that the immune response is still highly active at day 20 although the number of CAR-T cells decreased (Fig. 4I).

**Fig. 4.**
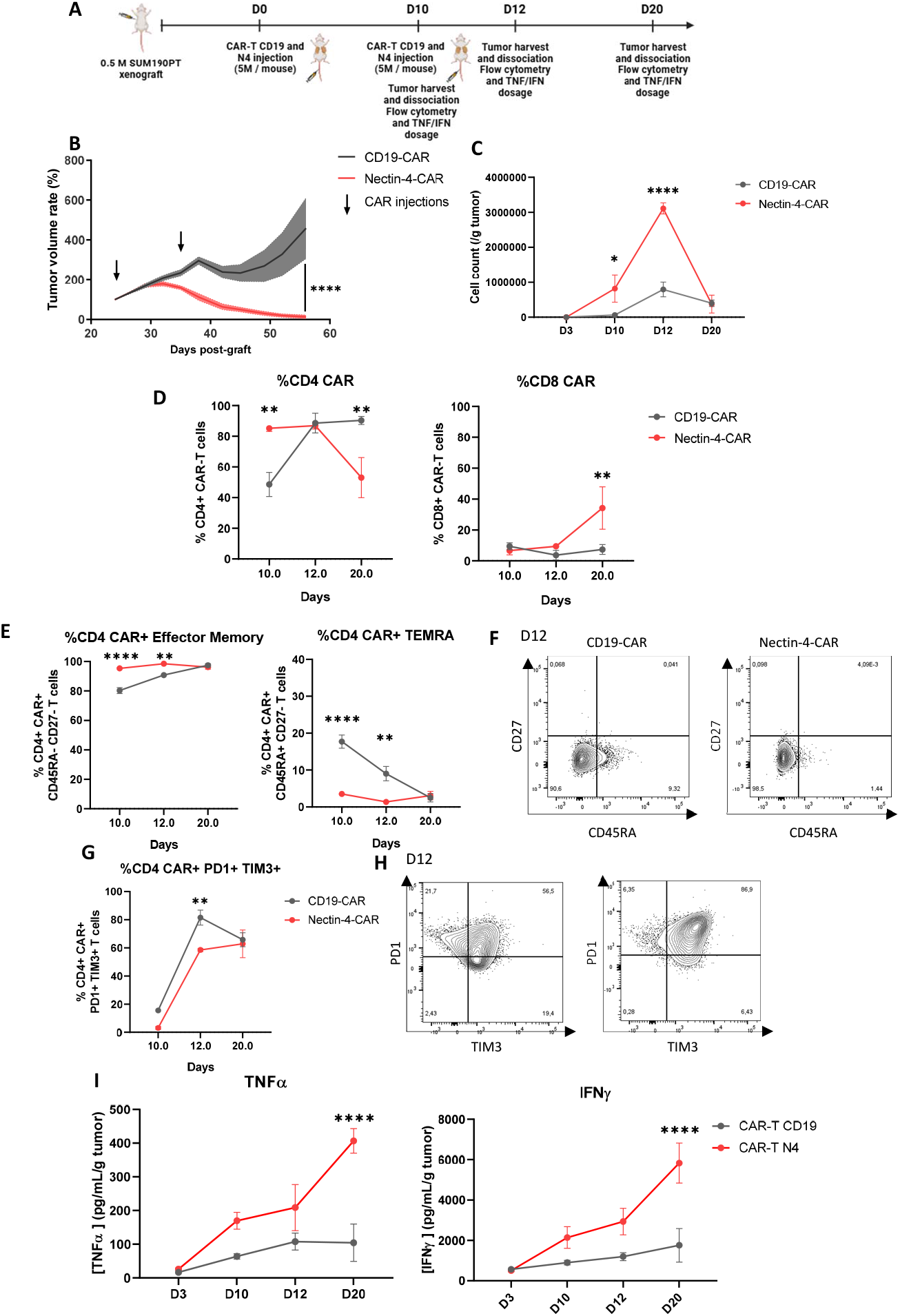
Mice double treatment with CARTN4 cells leads to tumor complete regression in human tumor xenograft models. A. Schematic illustration of the protocol. B. SUM190PT tumor total regression with a double injection of CARTN4 cells *in vivo*. SUM190PT cells were orthotopically injected into the mammary fat pad of female NSG mice. Mice received a double i.v. injection of CART19 as vehicle control or Nectin-4-CAR T cells. n = 5 CD19-CAR and n = 5 Nectin-4-CAR animals with 2 tumors each were examined in one experiment. C. CD3+GFP+ absolute count per gram of tumor is higher in Nectin-4 tumor until the 12^th^ day. Tumors were harvested and dissociated at day 10, 12 and 20 after CAR-T cells treatment. D. CD4 and CD8 rate in CAR-T cells showing a increase in the CD8 population at day 20 for CARTN4 cells. E. Effector memory T-cells rate in CD4 T-cells increases over time until reaching that of CARTN4 cells at day 20 whereas EMRA T-cells rate decreases. F. Flow cytometry results illustrating the higher frequency of TEMRA in CART19 cells and the higher frequency of effector memory T-cells in Nectin-4-CAR-T cells at day 12. G. PD1 and TIM3 double positive T-cells rate in CD4 T-cells in tumors at day 10. E. Flow cytometry results illustrating the higher frequency of PD1 and TIM3 double positive T-cells in CART19 over time until reaching that of CART1N4 at day 20. H. ELISA assay dosing TNFα and IFNγ in tumor dissociation supernatants revealing higher concentrations of both cytokines in CARTN4 / tumors whatever the day. 4 tumors were harvested from 2 mice per condition. Data in panels B, C, D, E, G and I are presented as Mean +/- SEM. P-values were calculated using a 2way ANOVA. Source data are provided in the Source Data file.

### Enfortumab vedotin-resistant (EVr) cells are susceptible to N4CART cell killing

To emphasize the superior activity of our N4CART cells, we decided to validate their killing potency using SUM190PT EVr resistant cells (SUM190PT EVr), derived from the sensitive SUM190PT cells ^18^. Both cells have comparable expression level of Nectin-4 as illustrated by flow cytometry (Fig 5A). SUM190PT EVr are resistant to EV showing unchanged viability when compared to an irrelevant CD30-ADC (Fig.5B). We next evaluated the efficacy of the N4CART cells and 19CART cells on SUM190PT EVr by co-culture experiments. Interestingly, although resistant to EV, SUM190PT EVr co-culture with N4CART cells generated CD4+ and CD8+ CAR+ T cells characterized by TNF-α and IFN-γ expression, equivalent to a co-culture with SUMP190PT (Fig 5C). Moreover, N4CART cells killed similarly SUMP190PT and SUM190PT EVr tumor cells as detected by caspase 3/7 in cytometry (Fig 5D-E). The co-culture with 19CART cells did not show any cytokine expression response (Fig 5C) and exhibited a decreased cytotoxicity when compared to N4CART cells (Fig 5D). We then, decided to test *in vivo* the tumor growth of SUM190PT EVr treated with N4CART cells or 19CARTcells (Fig 5F). We observed an important tumor regression when N4CART cells were injected twice and even observed 8/9 complete responses (one mouse has been stopped at day 73 for further IHC) analysis. As expected, the tumor growth was not controlled when 19CART cells were injected. These results illustrate the added value in using CAR-T cell immunotherapy in EVr tumors as compared to the current available treatments.

**Fig. 5.**
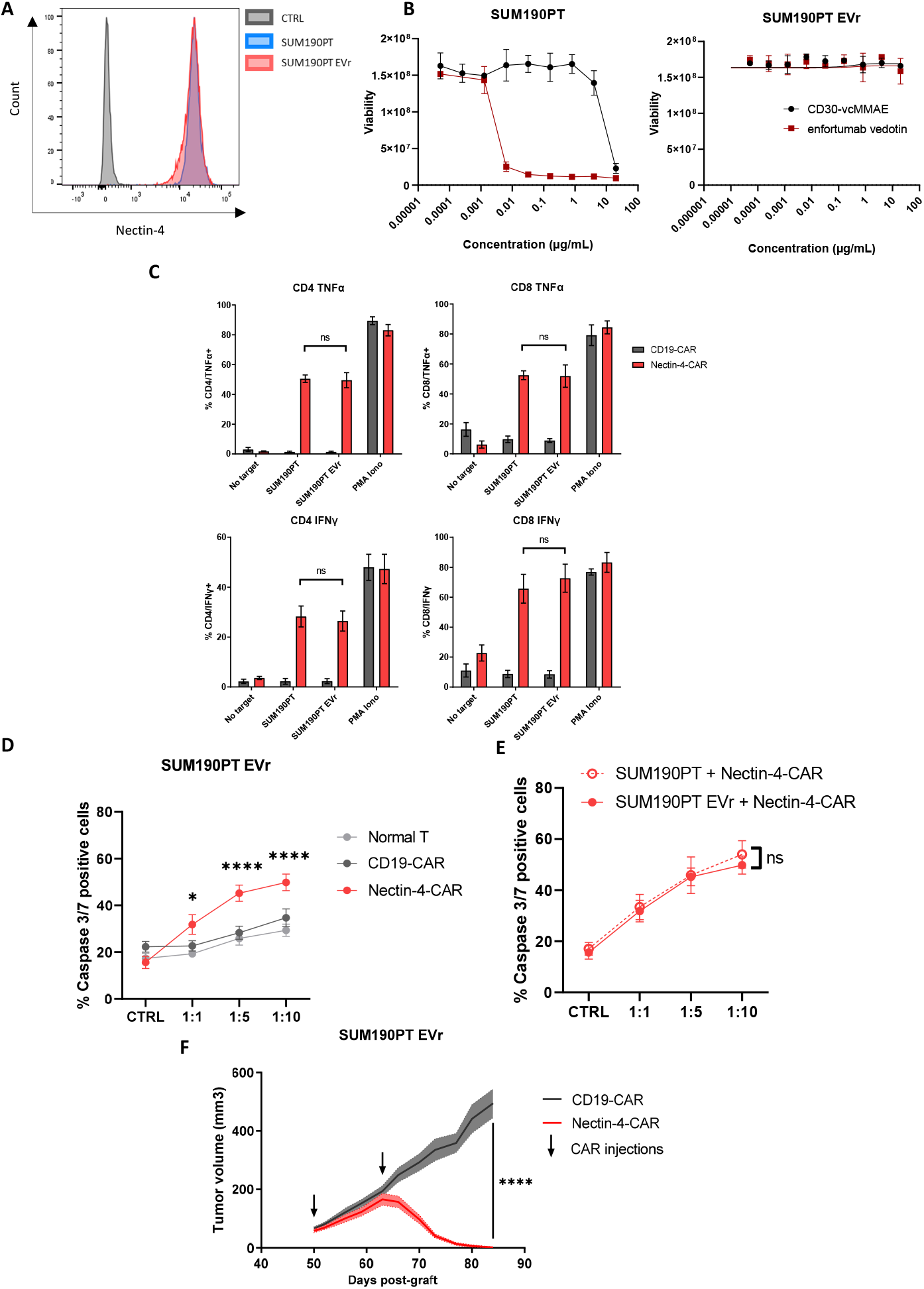
CARTN4 kill N4-positive tumor cell line which acquired a resistance to enfortumab vedotin after a long-term treatment with this ADC. A. Nectin-4 expression is similar on the EV sensitive and enfortumab vedotin resistant SUM190PT cell line. B. The EV sensitive and resistant SUM190PT cell line were treated with CD30-MMAE(negative control) and enfortumab vedotin. The SUM190 cell line was sensitive to enfortumab vedotin unlike the resistant cell line SUM190PT Evr. n = 2 biologic replicates were examined in one experiment. C. 4-hours TNFa and IFNg production assay showing an equal rate of CD4 and CD8 CAR-T cells producing cytokines in contact with SUM190PT and SUM190PT EVr. CD4 and CD8 T-cells are permeabilized and stained with anti-TNFa and IFNg antibodies. n=4 healthy donors were examined in two independent experiments. D. 4-hour caspase 3/7 cytotoxicity assay indicating a higher tumor killing of CARTN4 cells against SUM190PT resistant to enfortumab vedotin. n=9 healthy donors were examined in 5 independent experiments. E. In contact with SUM190PT or SUM190PT Evr CARTN4 cells have a comparable cytotoxicity effect. n=9 healthy donors were examined in 5 independent experiments. F. SUM190PT Evr tumor total regression with a double injection of CARTN4 cells *in vivo*. SUM190PT Evr cells were orthotopically injected into the mammary fat pad of female NSG mice. Mice received a double i.v. injection of CART19 as vehicle control or CARTN4 cells. n = 5 CD19-CAR and n = 5 Nectin-4-CAR animals with 2 tumors each. Data in panels B, C, D, E, and F are presented as Mean +/- SEM. P-values were calculated using a 2way ANOVA. Source data are provided in the Source Data file.

### Cancers models expressing Nectin-4 are susceptible to N4CART-killing

Interestingly, we tested the expression of Nectin-4 on different cancer cell models, a breast cancer model MCF7, a colorectal cancer HT-29, and a human bladder carcinoma RT112. The three cell models showed detectable Nectin-4 expression although the Nectin-4 expression level (MFI) tended to be low when compared to SUM190PT cells (Fig 6A). All cell models were confirmed to be resistant to EV (Fig 6B). In co-culture experiments of MCF-7 and RT112 cells with N4CART cells, these still showed increased killing. This was especially pronounced for RT112 and statistically relevant at higher E:T ratios for MCF7 cells (Fig 6C). HT-29 were modestly sensitive to the co-culture with N4CART cells although they were still more sensitive than to 19CART cells or regular T cells co-culture. These results show that EVr cells are still susceptible to N4CART cells killing *in vitro* independently of the type of tumor.

**Fig. 6.**
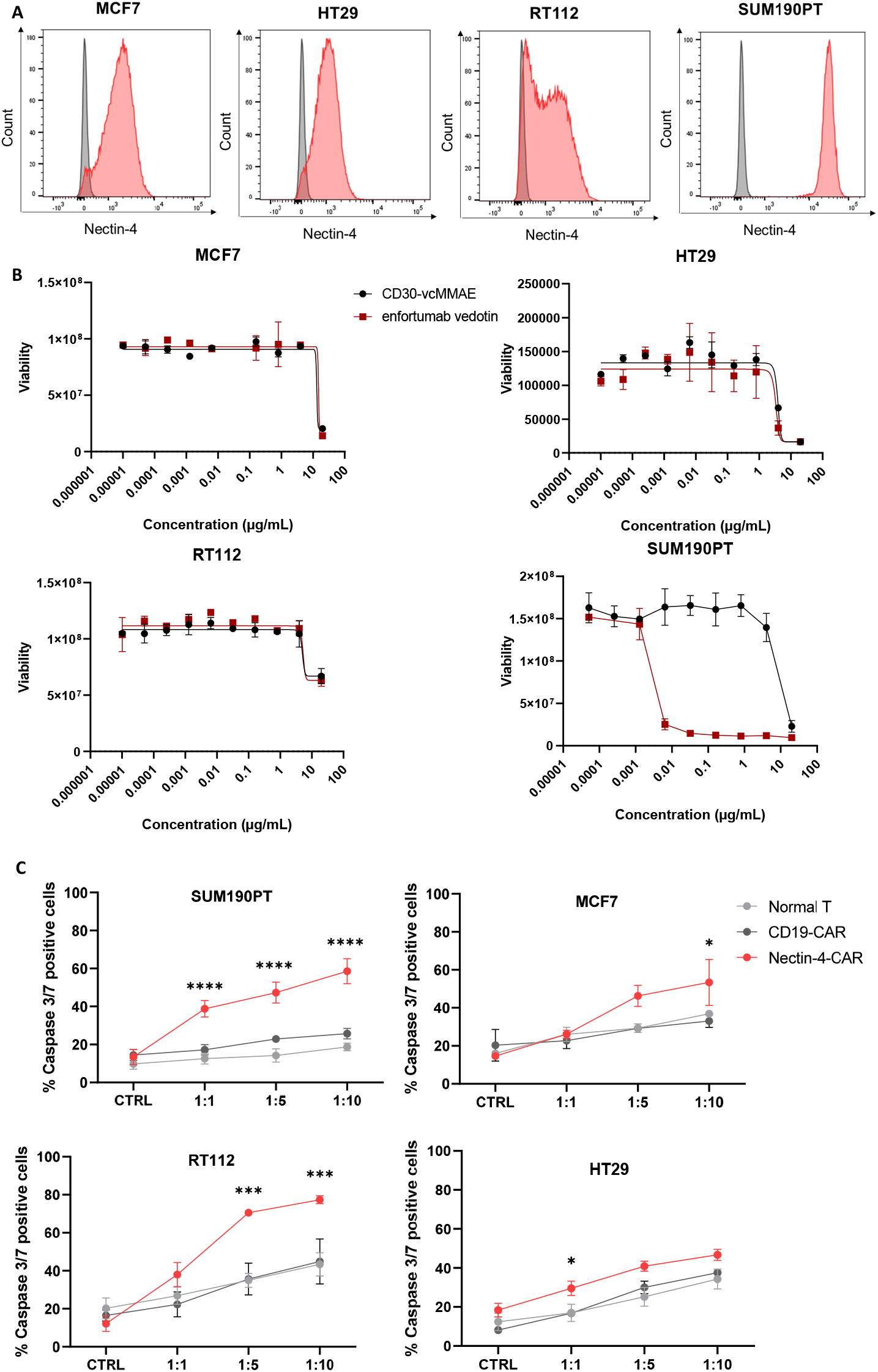
CARTN4 cells kill N4-positive tumor cell line which are naturally resistant to enfortumab vedotin. A. Nectin-4 expression on three cell lines in comparison with SUM190PT. B. MCF7, HT29, RT112 and SUM190PT cell lines were treated with CD30-MMAE(negative control) and enfortumab vedotin. MCF7, HT29 and RT112 cell lines are resistant to enfortumab vedotin whereas SUM190PT is sensible. n = 2 biologic replicates were examined in one experiment. C . 4-hour caspase 3/7 cytotoxicity assay indicating a tumor killing of CARTN4 cells against MCF7, HT29, RT112. n=4 healthy donors were examined in three independent experiments. Data in panels B, and C are presented as Mean +/- SEM. P-values were calculated using a 2way ANOVA. Source data are provided in the Source Data file.

### Improved N4CART transduction using baboon envelope pseudotyped LVs leads to efficient killing of TNBC PDX with low-level Nectin-4 expression

We recently demonstrated, the importance to use the best adapted lentiviral envelope pseudotype for cells that are difficult to transduce ^21^. Indeed, we showed that human NK cells were efficiently transduced using lentivirus bearing the BaEV envelope, this improved as well γδ T cells transduction ^22,23^. The use of the BaEV envelope compared to VSV-G allowed high-level expression of either Nectin-4 or CD19 CAR in human primary T cells (Fig 7A, B). To be closer from Nectin-4 expressed in patients tumors, these Nectin-4 or control CD19 CAR T cells were evaluated for their anti-cancer effect on patient derived xenograft (PDX) PDX317 tumors that we showed previously express intermediate level of Nectin-4 (HS:160) ^9^. NSG mice were inoculated with 0.25 × 10^6^ PDX317 cells in both flanks of the mammary fat pads. When tumors reached an average volume of 80-100 mm^3^, mice were randomized and received an i.v. adoptive transfer of 2.5 × 10^6^ CAR-T cells followed by a second identical dose of CAR T cells 10 days later. The injection of N4CART cells significantly decreased the tumor growth of PDX317 compared to the mice injected with 19CART cells (Fig 7C). Moreover, the clear infiltration of CD3+ T cells at day 13 in the tumor in vivo was only achieved upon injection of N4CART cells and nearly undetectable upon 19CART cell injection (Fig 7D). This last result validates the use and efficacy of N4CART cells, even in the context of PDX expressing a low amount of Nectin-4 under the condition that the CAR is strongly expressed.

**Fig. 7.**
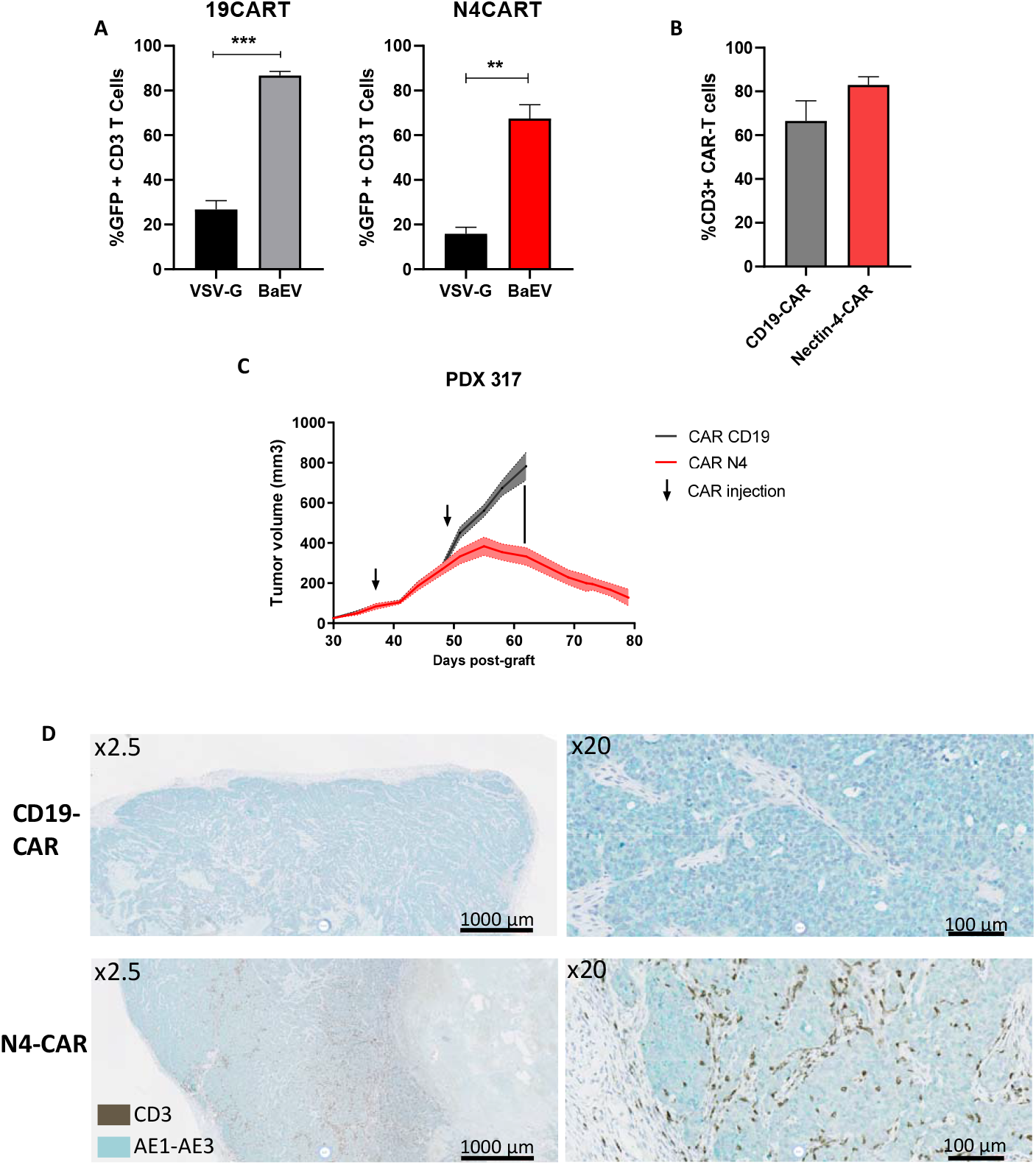
Mice treatment with baboon envelope pseudotyped lentiviral vectors to express Nectin-4 CAR cells leads to optimized expression and tumor growth control in N4-positive patient-derived xenograft models. A. CD19 and Nectin-4 CAR expression reported by the GFP expression in activated human T cells at Day 8 transduced at day 2 with either VSV-G or BAEV-TR lentiviral envelop N=3 healthy donors B. CD19 and Nectin-4 CAR expression reported by the GFP expression in activated human T cells after BAEV-TR pseudotyped lentiviral transduction. n=4 healthy donors were examined in two independent experiments. B. Anti-Nectin-4 IHC on the PDX 317 showing a low expression of Nectin-4. C. Low-positive N4 patient-derived xenograft growth control with a double injection of CARTN4 cells *in vivo*. PDX 317 were orthotopically injected into the mammary fat pad of female NSG mice. Mice received a double i.v. injection of CART19 as vehicle control or CARTN4 T cells. . n = 5 CD19-CAR and n = 5 Nectin-4-CAR animals with 2 tumors each. D. IHC on mouse SUM190PT tumor 13 days after CAR treatment showing a CARTN4 cell infiltration. Data in panels A, and C are presented as Mean +/- SEM. P-values were calculated using a 2way ANOVA. Source data are provided in the Source Data file.

## Discussion

Here, we described the use of a new Nectin-4 directed CAR. This nectin-4 CAR T cells showed efficiency against Nectin-4 breast cancer cell models either resistant or not to EV. Likewise, we showed *in vitro* that other cancer types naturally resistant to EV were susceptible to N4CART cell killing. Finally, we showed that N4CART cells exhibited *in vivo* efficacy in a TNBC PDX tumor mouse model expressing intermediate Nectin-4 levels.

The use of EV in clinical trials triggered several secondary effects including skin reactions, peripheral neuropathy and/or decreased white blood cell counts in mUC. A previous Nectin-4 CAR was engineered showing impressive response targeting MC38 colon cancer cells *in vivo* in an immune competent model ^20^. Authors, subsequently developed the hu-Nectin4-7.19 CAR T, a 4^th^ generation CAR construct that showed promising *in vitro* results and encouraging *in vivo* efficacy in preclinical tumor bearing NSG mice^21^. But unfortunately, in a phase I study NCT03932565 aiming to examine the safety and feasibility of Nectin4-7.19 CAR-T cell injections into patients revealed some major toxic side effects such as haemorrhagic rash and rash desquamation ^20^.

Since one of the most observed adverse events with the use of EV-ADC or previous CAR-T are skin reactions, we selected a scFv sequence that binds mouse Nectin-4 antigens but not human skin (Fig1). We selected a clone that could especially bind the mouse skin to address the eventual adverse effects on the skin of N4CART cells during our pre-clinical experiments (Fig 1C). Our results strengthen the need to use an appropriate antibody to target Nectin-4 that might indeed avoid or limit any secondary effects when developing an ADC or a CAR construct. We observed slight ventral hair loss in two mice out of 20 treated rat day 73 after initial treatment with N4CART cell (from experiments fig 5F). These mice were responding to the treatment and did not show any other adverse events. We analysed ventral skin of these mice and found Nectin-4 CAR T cells infiltration mostly localized in the basal epidermis and in the hair follicules in the dermis (Fig sup 7A). These data are in line with the reactivity found with the N4.78.6 antibody in mouse skin by IHC (Fig 1C) and may predict low-toxicity in preclinical and clinical settings^9^ .The N4.78.6 antibody was also selected based on the fact that it does not react with Nectin-4 expressed in human primary skin keratinocytes (NHEK) and in human and monkey skin tissue (Fig 1D-E). This property contrasts with EV-ADC known to bind skin and to induce skin toxicity (Fig 1D). We have recently produced an anti-Nectin-4 ADC, currently in Phase I study, whose antibody, 15A7.5, binds Nectin-4 with low affinity in human keratinocytes and human skin ^19^ . These antibodies recognized tumor Specific Antigenic Epitope (TSAE) but the mechanism of action is not yet completely elucidated. We cannot exclude that N4.78.6 differential binding between human and mouse skin tissue is related to the difference in apparent affinity observed between human Nectin-4 (12 nM) and mouse Nectin-4 (2 nM) (Fig 1B). However, we did not detect any staining in human skin with N4.78.6 concentration up to 10 mg/ml using a very sensitive detection method (Fig 1D). N4.78.6 binds to IgV distal domain of Nectin-4 but the exact location of the targeted epitope is not yet depicted.

One of the important aspects that we showed in here, is that, even Nectin-4 low expressing tumor cells (in the case of the PDX) were sensitive to N4CART cells killing. There have been several studies in the field showing that one of the problems of CAR-T cells, and especially CD19 CAR-T cells, is their ability to recognize targets that only express a high amount the antigens ^24,25^ and might thus be responsible of relapse in low-antigen expressing patients ^26^. Indeed, Nectin-4 protein expression is low to moderate in different carcinomas and in paired primary vs metastatic urothelial cancer (mUC). These data confirm the need of an appropriate therapy to target these low expressing tumors ^27^. Some CAR-contruct engineering already exists to overcome this potential resistance with the use of the HLA-independent T cell receptors (HIT receptors), which are highly sensitive to low antigen expressing tumors ^28^ . Additionally, new signaling modules have been proposed such has logic-gated intracellular network (LINK) CAR constructs to decrease the activation threshold upon antigen encountering ^29^. In the case of N4CART cells presented here, more effort should be made to fully answer this aspect, but our preliminary observations are promising since N4CART cells recognize already cells with low target antigen expression such as the PDX317 tumor cells presented here.

Interestingly, we also showed an *in vitro* efficacy of N4CART cells against low-Nectin-4 expressing colorectal cancer HT-29 cells and human bladder carcinoma RT112. Indeed, EV is already approved in clinic in the context of advanced or metastatic urothelial carcinoma ^16^. Although our results focus mainly on TNBCs, future work is needed to test the *in vivo* efficacy of N4CART cells against different types of cancers.

Thus, N4CART cells that we presented here, already demonstrate an added value to the use of EV or other developed Nectin-4 CAR-T cells, making it a new safe alternative that might be of great benefit for patients with tumors expressing Nectin-4. Indeed, combinations with immune checkpoint inhibitors such as with anti-PD-1 antibodies and EV have been initiated in the clinic showing promising results, especially in bladder cancer ^30^. Such a strategy can also be considered with the use of N4CART cells to further enhance their anti-tumoral activity.

Here, we are presenting a 2^nd^ CAR generation construct. However, numerous approaches have been used to enhance the potency of CAR-T cells by “arming them” in many different ways. Several modules might be added to our present CAR such as the co-expression of cytokines (i.e. IL-12, IL-15, IL-21 …) ^31^, recently a hu19CART-IL-18 module was added in a clinical trial showing feasibility and promising results in patients with relapsed or refractory lymphoma ^32^, the expression of homing receptors such as CXCR6 ^33^ or the modification directly of the ζ-chains (1XX CAR T cells) ^34^ or co-stimulation modules ^35^. This, to diminish the exhaustion phenotype, improve the persistence and the CAR signaling. Another important aspect in the future would be to control the virus insertion to limit the off-target effects to a precise locus ^36^. Finally, we also suggested the genetic targeting of intracellular inhibiting proteins that might be of interest in combination with such a CAR ^37^. However, the choice of “weapons” might be highly dependent of the type of cancer that is targeted and shall be carefully explored in the future.

Finally, we previously showed that the use of baboon envelope pseudotyped lentiviral vectors was efficient to genetically modify difficult to transduce cells such as primary human NK cells or human mammary epithelial cells ^21,23,38^ . In here, the use of baboon envelope pseudotyped lentiviral vectors to express Nectin-4 CAR in αβ T cells showed an impressive high level CAR expression coupled to a great anti-tumoral response with the PDX317. Although high transduction efficiency and integration of the transgene is important when CAR-T cells are produced for the patient in clinical settings, sometimes less than 10% CAR T cells are positive during the cell production for clinic, this jeopardizes the efficiency or the use of the cell product in clinic ^39^. The use of this new BAEV envelope is thus an interesting alternative to augment even more transductions rates in αβ T cells while producing more efficient CAR-T cells for clinical applications.

Altogether, we describe a new Nectin-4 CAR-T construct that is efficiently targeting Nectin-4 low-expressing tumors without triggering any adverse effects. Although more efforts are required to ensure the safety and improve even more this construct in the future, the Nectin-4 CAR T presented here already represents a promising tool to be developed in clinic in the future.

## Materials and Methods

### Antibodies and reagents

Monoclonal antibodies (mAbs) directed against Nectin-4 were produced after mice immunisation with human recombinant dimeric extracellular Nectin-4 protein fused to human IgG1 Fc fragment. Screening was performed on transfected vs non transfected Nectin-4 Cos cells. N4.78.6 and N4.72.1 mabs were selected in this study (**Fig.1**). Enfortumab vedotin was purchased to Evidentics GmbH (Germany).

### Cells and cell lines

The SUM190 cancer cell line expressing high level of Nectin-4 was from two origins: One kindly provided by Dr Ethier (Karmanos Cancer Center, Detroit, MI, USA) and was cultured in Ham’s F12 medium with 2% FBS, non-essential amino acids, 10 μg/mL insulin, 1 μg/mL streptomycin and 2 mM glutamine; the other purchased from BioIvt (Germany) was cultured in the same condition supplemented with 6.7 ng/mL Triiodo-L-tyronine. Absence of mycoplasma contamination was regularly controlled by PCR (Eurofins Genomics, Germany), and MycoAlert™ PLUS test (Lonza, Switzerland). Both cell lines expressed similar cell surface levels of Nectin-4. MCF7 and HT-29 cell lines for this study were purchased from American Type Culture Collection (ATCC,Manassas, VA 20110, USA). RT112 cell line was purchased from DSMZ Cell Dive (Braunschweig, Germany). Cell lines were cultured as instructed. Cell line authentication tests were done by Eurofins Genomics.

### Transfection experiments

COS cells grown to 50–80% confluency were transfected with the appropriate cDNA expression plasmids by using the FuGENE™6 reagent method. The cells were cultivated for 1 day, and the medium was replaced. Cells were directly processed in the case of transient expression assays. Vectors used in this study are: 1-p3XFLR4.C1 - Human cDNA Nectin-4/*PVRL4* (vector p3XFLAG) ^7^. 2-PVRL4_ Omu23587 – Mus musculus cDNA Nectin-4 / *PVRL4* (vector pcDNA3.1) (Genscript, NJ, USA).

### Construction of chimeric antigen receptor and lentiviral vector production

The variable region sequences of heavy (VH) and light chain (VL) of the anti-Nectin-4 N4.78.2 mAb were used to design scFv connected with a G_4_S linker (Patent EP25201525.0). The CD19-scFV was proposed by VectorBuilder. The second-generation CARs consist of a human CD8 leader, Nectin-4- or CD19-scFv, hinge and TM regions of the human CD8 molecule, 4-1BB intracellular domain sequence and CD3z intracellular domain sequence. The eGFP sequence is included via a T2A ribosomal skipping sequence in the construct to allow detection of transduced T cells. The lentiviral vector encoding the CARs was produced by VectorBuilder.

### Lentiviral vector production

The CAR sequences were then integrated into VSV-G pseudotyped third-generation lentivirus by VectorBuilder. BaEV-TR pseudotyped third-generation lentivirus were produced by transfection of HEK293T cell line with 3 plasmids: CMVBaevTR (patented and available from Els Verhoeyen), psPAX2 (Addgene), CAR encoding plasmid (VectorBuilder). HEK293T were co-transfected using transfection reagent Lipofectamine LTX (ThermoFisher Scientific). After 48 hrs, the supernatants were harvested and removed any cell debris by filtering through a 0.45 μm filter. Virus were then concentrated 100 times using Lenti-X Concentrator (Takara).

### T-cell transduction

Peripheral blood mononuclear cells (PBMCs) from healthy donors were isolated by gradient centrifugation at 800g for 30min using medium for separation of lymphocytes (Eurobio Scientific) at room temperature. Human PBMCs were activated with purified anti-human CD3 mAb ( clone OKT3, Miltenyi Biotec) at 50 ng/mL and purified anti-human CD28 mAb ( clone CD28.2, Biolegend) at 125 ng/mL in CTS Optimizer™ T-cell expansion media (ThermoFisher Scientific) supplemented with 5% human AB serum (Fisher Scientific), 5% CTS Immune cell serum (ThermoFisher Scientific) and 300 U/mL IL-2 (Miltenyi Biotec) at density of 2x10^6^ cells/mL for 48 hrs. Activated T cells were transduced with VSV-G envelope lentivirus at 0.5x10^6^ cells/mL at MOI 10 and spinoculated for 2h at 1200g at 32°C. T cells transduced with BAEV-TR envelope lentivirus were transduced at MOI 10 with Retronectin^®^ as manufacturer’s procedure. The transduced T cells were maintained in medium containing IL-2. Transduction efficiency was determined by analysing CAR-T cells GFP expression by flow cytometry. Activated but non-lentiviral transduced cells were included as normal T controls.

### Cytotoxicity assay

The antigen-specific tumor cell lysis by N4CART cells was determined by caspase 3/7 expression assay. Untransduced T cells, 19CART cells or N4CART cells were co-cultured with stained target cells with at E:T of 10:1, 5:1 and 1:1 ratio for 4 hrs at 37°C. The quantification of apoptotic cells was performed using the CellEvent™ Caspase-3/7 Green Flow Cytometry Assay Kit (Thermo Fisher Scientific) following manufacturer’s instructions.

### Cytokine Release Assays

Normal T, 19CART or N4CART cells were co-cultured with target cells at E:T of 5:1 ratio or with PMA (50 ng/mL) and ionomycin (2 µg/mL) for 4 hrs at 37°C. Cell-free supernatants were harvested for testing TNFα and IFN-γ secretion by ELISA kits (BD Biosciences) according to the manufacturer’s instructions.

### Analysis of cytokine production

For the measurement of intracellular TNFα and IFN-γ production, normal T, 19CART or N4CART cells were co-cultured with target cells at E:T of 5:1 ratio or with PMA (50 ng/mL) and ionomycin (2 µg/mL) with Monensin and Brefeldin A (BD GolgiPlug™ and GolgiStop™) for 4 hrs at 37°C. CAR-T cells were subjected to surface staining and intracellular staining to detect TNFα and IFNγ production by flow cytometry.

### Flow cytometry

Cells (10,000 – 50,000) expressing Nectin-4 (naturally or transfected) were incubated with indicated antibodies. After washing, cells were then stained with phycoerythrin-conjugated goat anti human antibody (5 µg/mL, Jackson ImmunoResearch). After fixation, cells were stained with a viability dye (e780, Invitrogen) before flow cytometry acquisition and analysis. CAR expression on T cells was detected by PE-Vio770-conjugated CD3 (clone BW264/56, Miltenyi Biotec), GFP expression and biotin-conjugated Nectin-4 recombinant protein and streptavidin-APC (BioLegend). CAR-T cells were phenoyped with a panel of monoclonal anti-human antibodies as follows: PE-Vio770-conjugated CD3 (clone BW264/56, Miltenyi Biotec), VioBlue-conjugated CD4 (clone VIT4, Miltenyi Biotec), VioGreen-conjugated CD8 (clone BW135/80, Miltenyi Biotec), APC-conjugated CD45RA (clone T6D11, Miltenyi Biotec) and PE-conjugated CD27 (clone M-T271, Miltenyi Biotec). Cytokines in CAR-T cells were detected by APC-conjugated TNFα (clone cA2, Miltenyi Biotec) and PE-conjugated IFNγ (clone 45-15, Miltenyi Biotec). In most assays, cells were stained with LIVE/DEAD™ Fixable Near IR (780) Viability Kit (ThermoFisher Scientific) to exclude dead cells from analysis. Flow cytometry data were acquired with a FACSCanto™ system (BD Biosciences) using DIVA software according to the manufacturers’ instructions. Data were analyzed using FlowJo software.

### *In vivo* studies in mice

All experiments were done in agreement with the French Guidelines for animal handling and approved by the local ethics committee (CRCM for SUM190 PT, PDX TNBC317 APAFiS#13349, 16487 and 21988). During the study, the care and use of animals was conducted in accordance with the regulations of the Association for Assessment and Accreditation of Laboratory Animal Care (AAALAC). Female NOD/SCID/γc null (NSG) and male NMRI-nude mice were obtained from Charles River laboratories. Mice were housed under sterile conditions with sterilized food and water provided *ad libitum* and maintained on a 12-h light and 12-h dark cycle. In NSG mice, cells and PDX were inoculated in both flanks in the mammary fat pads with 0.25 × 10^6^ cells (except for experiment fig 3 0.5 × 10^6^ cells) suspended in 50% phenol red-free Matrigel (Becton-Dickinson Bioscience). Mice were treated when tumor reached average volume of 50-150 mm^3^. Mice were treated i.v. with 5 × 10^6^ cells CAR-T cells (2.5 × 10^6^ cells for PDX317 experiment Fig 7) with IL15 2.5µg (BioTechne #247-ILB-025) /IL15R (BioTechne #7194-IR-050) 10µg. Tumor growth was monitored by measuring with a digital caliper and by calculating the tumor volume (length × width^2^ × height × π/6). All animals were randomly assigned into treatment groups (N=5 with 2 tumors injected each flank i.e 10 tumors for each group), such that the mean and the median tumor volume for each group were comparable. Animal weight was monitored every 3-4 days to evaluate the toxicity of the different treatments. Mouse weight loss >20%, total tumor volume >1500 mm^3^, ruffled coat and hunched back, weakness, and reduced motility were monitored and considered as endpoints.

### IHC staining

Optimal cutting temperature (OCT) medium frozen blocks were mounted on disks and cryosection were performed at -20°C on a Microm cryostat (NX70). Cryosections (5 μm) were mounted on superfrost + slides (VWR). Sections were arranged on the slide and fixed in methanol at -20°C for 5 minutes. Immunohistochemistry was performed on the Ventana Discovery Ultra plateform. Endogenous peroxidases were inhibited by immersing the slides in hydrogen peroxide (H_2_O_2_) as part of Roche ChromoMAP DAB kit protocol. Concentration of 10 µg/mL of mouse N4.78.6 and N4.72.1 mabs were determined as optimal. To mitigate the non-specific binding of the secondary antibody to the mouse tissue, a Rabbit anti-mouse IgG (4 µg/mL, Abcam) was added following primary Ab incubation. The Multimer Omni-MAP anti-Rabbit HRP or the Multimer anti-Rabbit HQ and the anti-HQ HRP (Roche Diagnostic) was then used with ChromoMAP DAB to perform the DAB staining according to manufacturer’s instruction (Ventana Discovery Ultra automat). Finally, the counterstaining was done with hematoxylin and slides were cleaning, deshydrated and coverslip added with permanent mounting media. An assessment of both the staining intensity and the proportion of stained cells was performed. Immunostained slides were digitalized with a whole slide scanner (Nanozoomer 2.0HT, Hamamatsu Photonics, Hamamatsu, Japan) at 40X. The resulting ndpi image files were analyzed using CaloPix Research software (TRIBVN Healthcare, Paris, France).

IHC experiments on FFPE embedded tissues 3µm sections were performed by multiplex immunohistochemistry (mIHC) on the Ventana Discovery Ultra platform (Roche Diagnostic). Anti-CD3e (Clone 2GV6, Roche), anti-GFP (EPR14104-89, Abcam) and anti AE1-AE3 (Roche) were used at recommended by the manufacturer. After deparaffinization, antigens retrieval was performed with Disco CC1 buffer pH 8-8.5. The primary antibodies were incubated sequentially : first, rabbit anti GFP antibody 1,86µg/ml was incubated followed Multimer anti Rabbit HQ then an anti-HQ HRP detection with the Purple substrate; second, mouse anti human cytokeratin 0,75µg/ml was incubated followed by a linker rabbit anti mouse IgG to mitigate the non-specific binding of the Roche multimer to the mouse tissue then OmniMAP anti-Rabbit HRP was added with the Teal substrate; third, rabbit anti human CD3e syringe RTU 0,4 ug /ml was incubated followed by Multimer anti Rabbit HQ then an anti-HQ HRP detection with the Yellow substrat. Counterstaining was done with hematoxylin and slides were cleaning, dehydrated and coverslip added with permanent mounting media. Immunostained slides were digitalized with a whole slide scanner (Nanozoomer 2.0HT, Hamamatsu Photonics, Hamamatsu, Japan) at 40X. The resulting ndpi image files were analyzed using CaloPix Research software (TRIBVN Healthcare, Paris, France).

### Statistical analyses

Prism software (GraphPad) was used for statistical analyses. Nonparametric tests were used according to the comparison setting (Mann-Whitney or Kruskal-Wallis and Dunn multiple comparison). To compare tumor sizes between treatment groups, 2-way ANOVA test and Bonferroni multiple comparison tests were used.

## Data availability

The data generated in this study are available upon request from the corresponding author

