## supplemental Figures for "CAR T cells targeting Nectin-4 safely overcome resistance to anti-Nectin-4 antibody-drug conjugate in solid tumors"

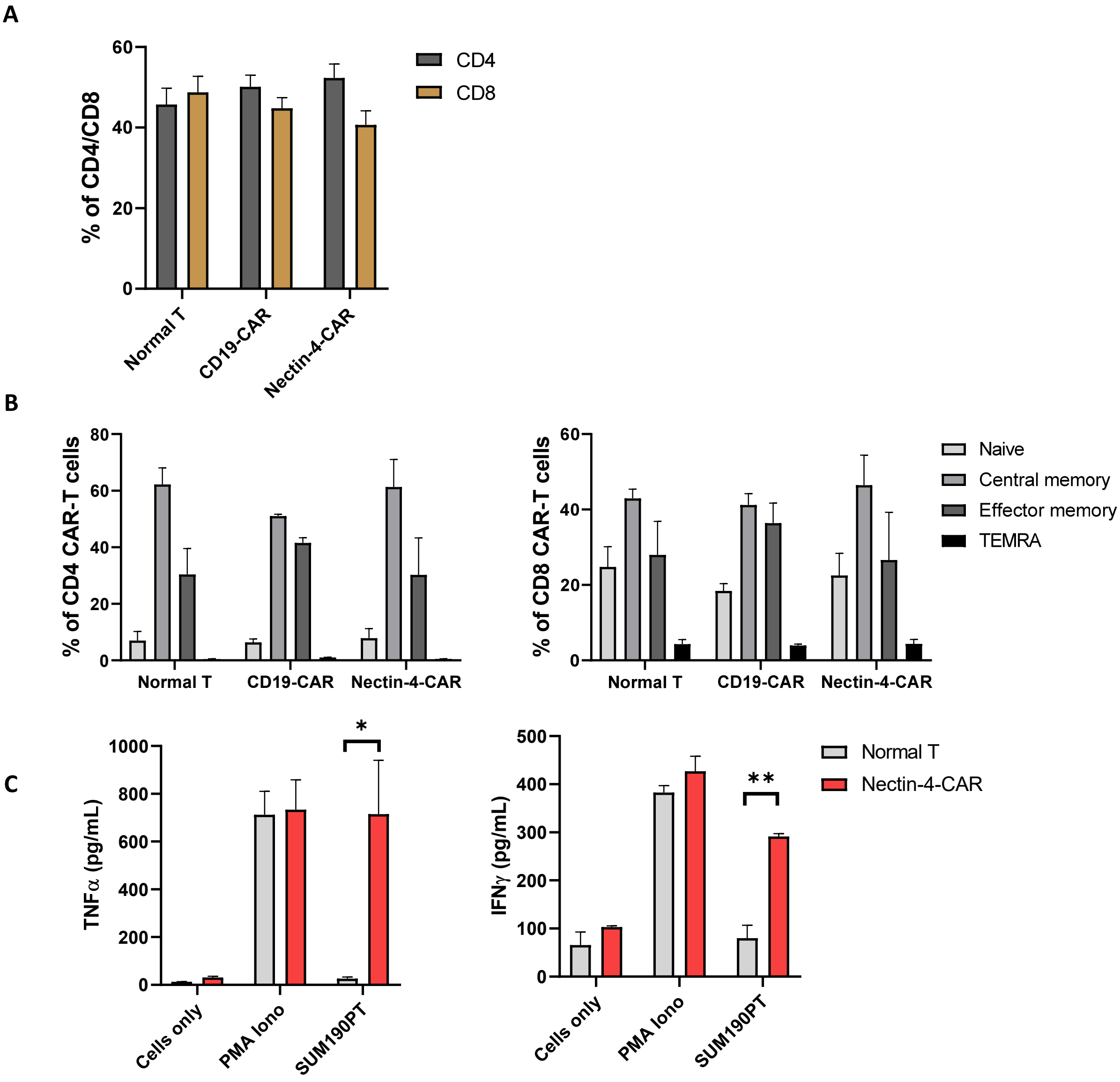


**Supplementary Fig. 1. Nectin-4-CAR generation**

A. CD4 and CD8 proportion are about 50% and are the same between CAR conditions. B. Differentiation state of CAR-T cells C. ELISA assay dosing TNFα and IFNγ producted by anti-Nectin-4-CAR-T cells in contact with SUM190PT cell line.

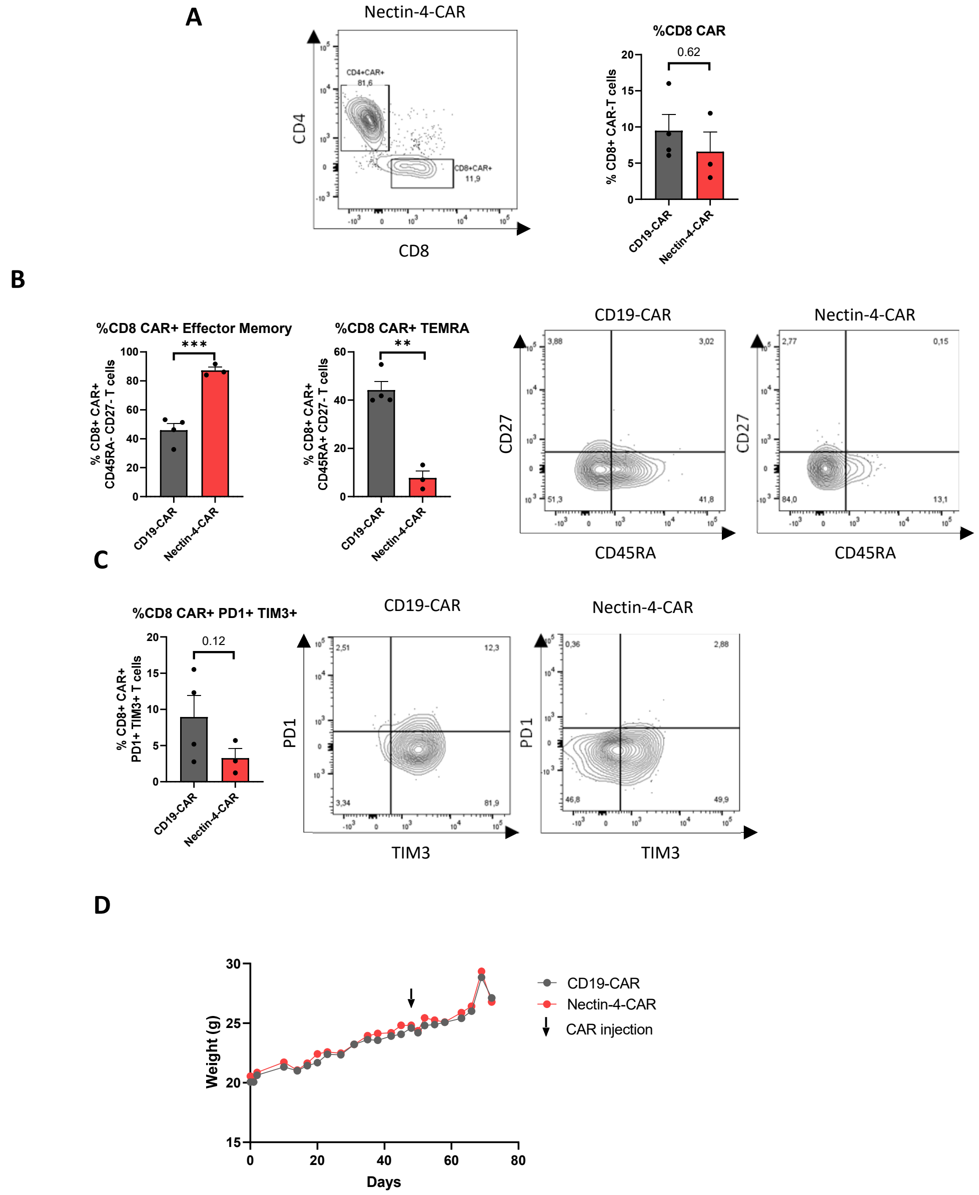


**Supplementary Fig. 2. CAR-T cells analysis after mice single treatment with N4-CAR-T cells.**
A. CD4 and CD8 proportion in Nectin-4-CAR-T cells at day 10. B. Flow cytometry results illustrating the CARTN4 and CART19 phenotype of CD8 T-cells in tumor at day 10 C. PD1 and TIM3 double positive T-cells rate in CD8 T-cells in tumors at day 10.D. Weight of mice treated with Nectin-4 and CD19-CAR-T cells showing that all mice have the same weight evolution. No toxicity is detectable in the weight variable.

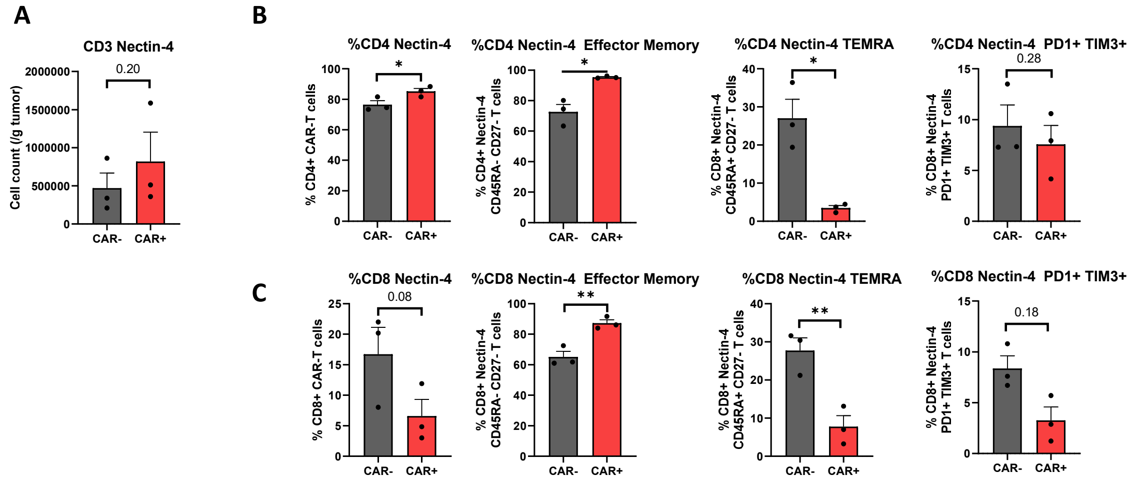


**Supplementary Fig. 3. Positive and negative CAR-T cells comparison after mice single treatment with N4-CAR-T cells.**

A. CD3 intratumoral infiltration status between untrasfected and CARTN4+ cells. B. CD4 T-cells phenotype comparison between positive and negative CARTN4 cells at day 10. C. CD8 T-cells phenotype comparison between positive and negative CARTN4 cells at day 10.

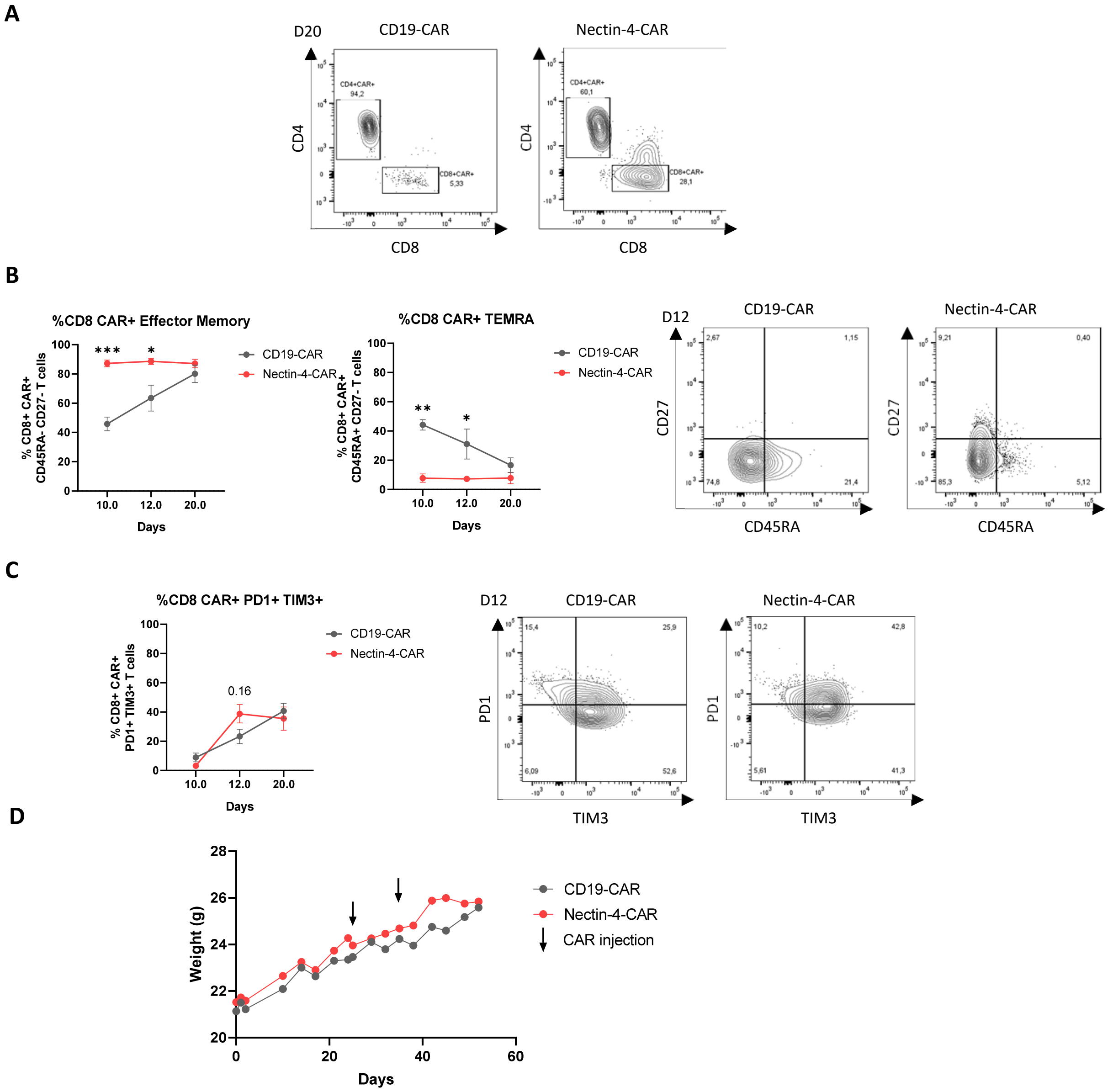


**Supplementary Fig. 4. CD8 CAR-T cells analysis after mice double treatment with N4-CAR-T cells.**

A. CD4 and CD8 proportion in CD19 and Nectin-4-CAR-T cells at day 20. B. Effector memory and EMRA T-cells rate in CD8 T-cells in tumors C. PD1 and TIM3 double positive T-cells rate in CD8 T-cells in tumors. D. Weight of mice treated with Nectin-4 and CD19-CAR-T.

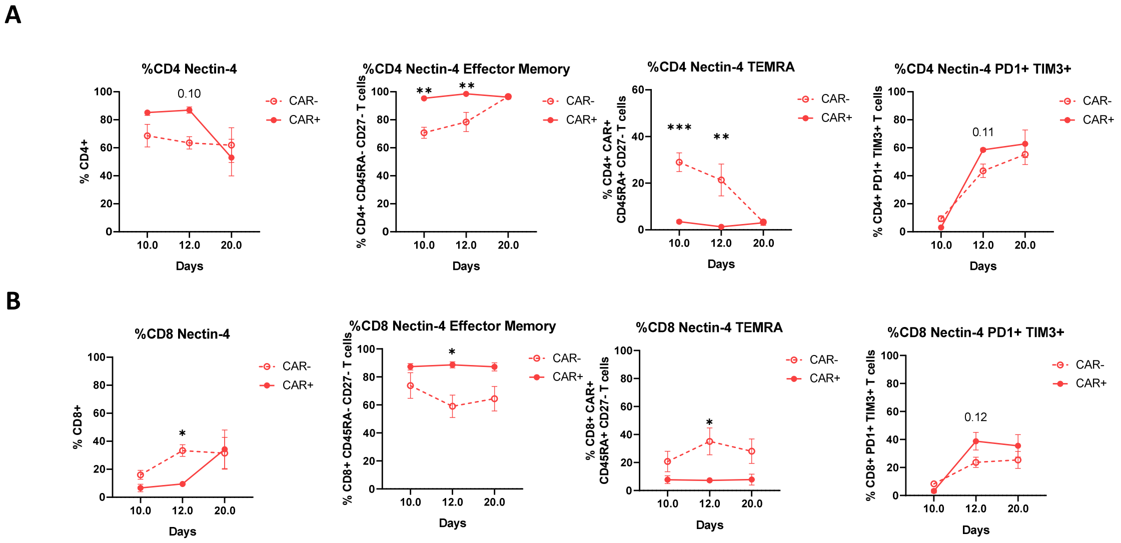


**Supplementary Fig. 5. Positive and negative CAR-T cells comparison after mice double treatment with N4-CAR-T cells.**

A. CD4 T-cells phenotype comparison between positive and negative CARTN4 cells. B. CD8 T-cells phenotype comparison between positive and negative CARTN4 cells.

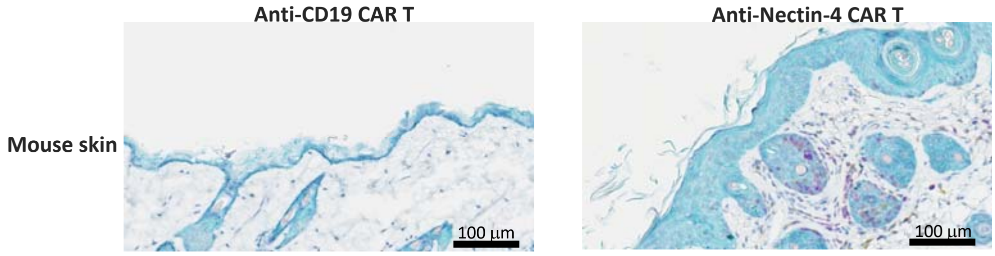


### Supplementary Fig. 6. Nectin-4-CAR T infiltration in mouse skin

IHC on mouse skin, 73 days after double CAR treatment showing a Nectin-4-CAR T infiltration in skin and hair follicles (experiment in Fig. 5F).
